# Genetic and pharmacological reduction of CDK14 mitigates α-synuclein pathology in human neurons and in rodent models of Parkinson’s disease

**DOI:** 10.1101/2022.05.02.490309

**Authors:** Jean-Louis A. Parmasad, Konrad M. Ricke, Morgan G. Stykel, Brodie Buchner-Duby, Benjamin Nguyen, Amanda Bruce, Haley M. Geertsma, Eric Lian, Nathalie A. Lengacher, Steve M. Callaghan, Alvin Joselin, Julianna J. Tomlinson, Michael G. Schlossmacher, William L. Stanford, Jiyan Ma, Patrik Brundin, Scott D. Ryan, Maxime W.C. Rousseaux

**Affiliations:** University of Ottawa Brain and Mind Research Institute, Ottawa, ON; Department of Cellular and Molecular Medicine, University of Ottawa, Ottawa, ON; Aligning Science Across Parkinson’s (ASAP) Collaborative Research Network, Chevy Chase, MD; Department of Molecular and Cellular Biology, University of Guelph, Guelph, ON; Program in Neuroscience, Ottawa Hospital Research Institute, Ottawa, ON; Ottawa Institute for Systems Biology, University of Ottawa, Ottawa, ON; Hotchkiss Brain Institute, Department of Clinical Neurosciences, University of Calgary, Calgary, AB; Parkinson’s Disease Center, Department of Neurodegenerative Science, Van Andel Institute, Grand Rapids, MI

**Keywords:** α-Synuclein, Parkinson’s disease, Cyclin-dependent kinase 14, Neurodegeneration, Therapeutics

## Abstract

Parkinsona’s disease (PD) is a debilitating neurodegenerative disease characterized by the loss of midbrain dopaminergic neurons (DaNs) and the abnormal accumulation of α-Synuclein (α-Syn) protein. Currently, no treatment can slow nor halt the progression of PD. Multiplications and mutations of the α-Syn gene (*SNCA*) cause PD-associated syndromes and animal models that overexpress α-Syn replicate several features of PD. Decreasing total α-Syn levels, therefore, is an attractive approach to slow down neurodegeneration in patients with synucleinopathy. We previously performed a genetic screen for modifiers of α-Syn levels and identified CDK14, a kinase of largely unknown function as a regulator of α-Syn. To test the potential therapeutic effects of CDK14 reduction in PD, we ablated Cdk14 in the α-Syn preformed fibrils (PFF)-induced PD mouse model. We found that loss of Cdk14 mitigates the grip strength deficit of PFF-treated mice and ameliorates PFF-induced cortical α-Syn pathology, indicated by reduced numbers of pS129 α-Syn-containing cells. In primary neurons, we found that Cdk14 depletion protects against the propagation of toxic α-Syn species. We further validated these findings on pS129 α-Syn levels in PD patient neurons. Finally, we leveraged the recent discovery of a covalent inhibitor of CDK14 to determine whether this target is pharmacologically tractable *in vitro* and *in vivo*. We found that CDK14 inhibition decreases total and pathologically aggregated α-Syn in human neurons, in PFF-challenged rat neurons and in the brains of α-Syn-humanized mice. In summary, we suggest that CDK14 represents a novel therapeutic target for PD-associated synucleinopathy.

## INTRODUCTION

Parkinson’s disease (PD) is a neurodegenerative disease that affects over 10 million individuals worldwide (1,2). Individuals with PD present with motor symptoms such as bradykinesia, rigidity, shuffling gait, and resting tremor, as well as non-motor symptoms such as constipation, anosmia and sleep disturbances (3–5). Neuropathologically, PD is characterized by the loss of dopaminergic neurons (DaNs) in the *substantia nigra pars compacta* (SN), as well as the accumulation of α-synuclein (α-Syn) containing inclusions, termed Lewy bodies and Lewy neurites (collectively: Lewy pathology) in surviving neurons (6–8). Current treatments address motor deficits, but are less effective on non-motor aspects of the disease and cannot slow down nor halt neurodegeneration in PD (9). Therefore, identifying new ‘druggable’ targets for PD is clearly warranted. In addition to being abundantly present in Lewy pathology, point mutations and multiplications in the gene encoding α-Syn, *SNCA*, underlie monogenic variants of PD (10,11). Increased levels of *SNCA* mRNA are also observed in laser-captured SN DaNs from PD patients (12), and animal models that overexpress α-Syn replicate several features of PD (13–15). Thus, there is a clear link between increased α-Syn dosage and PD pathogenesis, highlighting the crucial role of α-Syn in the manifestation of PD (16–19).

Since α-Syn dosage is linked to PD, decreasing total α-Syn levels may be a feasible approach to mitigate neurodegeneration in PD patients, regardless of whether oligomeric or fibrillar α-Syn is the toxic culprit. *Snca*-knockout (KO) mice are viable and fertile but display mild cognitive impairments, suggesting that a modest amount of cerebral α-Syn is required to accomplish its physiological role in the synapse (20,21). Titration of excessive α-Syn levels in a non-invasive manner would thus be beneficial in treating a chronic neurodegenerative disease like PD. Specifically, an orally available drug capable of mitigating α-Syn toxicity could offer a minimally invasive approach, a feature particularly important in treating a chronic illness. A pooled RNA interference screen investigating ‘druggable’ modifiers of α-Syn levels identified cyclin-dependent kinase 14 (CDK14, a.k.a. PFTK1; Cdk14 or Pftk1 in mice) as a regulator of α-Syn (22). CDK14 is a brain-expressed protein kinase with a largely unknown biological function (23). Its expression is upregulated in certain cancers, such as esophageal and colorectal cancer, for which it has generated attention as a therapeutic target (24,25). In fact, this has led to the recent development of FMF-04-159-2, a potent, covalent inhibitor of CDK14 (26).

Since decreasing CDK14 leads to a mild reduction in endogenous α-Syn levels (22), we hypothesize that genetic and pharmacological inhibition of CDK14 reduces α-Syn pathology and PD-like phenotypes in mice and human cells. To test this hypothesis, we examined the consequence of *Cdk14* reduction on PD-like features in the preformed α-Syn fibrils (PFFs)- induced PD mouse model. We further explored the role of CDK14 in mediating α-Syn spread in primary neurons. We also tested the effect of CRISPR/Cas9-mediated reduction of *CDK14* in human neurons carrying the PD-linked *SNCA* A53T mutation. Lastly, using the covalent CDK14 inhibitor, we investigated if pharmacological inhibition of CDK14 is sufficient to decrease α-Syn levels in rodent and human neurons. In summary, we show that decreasing CDK14, both genetically and pharmacologically, reduces α-Syn accumulation, spread, and modifies α-Syn aggregation.

## MATERIALS AND METHODS

### Mouse strains

*Cdk14^+/-^* mice were generated by the Gene Targeting and Transgenic Facility of Texas A&M Institute for Genomic Medicine (TIGM), on a mixed (129/SvEvBrd x C57BL/6) background (as described previously (27). *Cdk14*^+/-^ mice were backcrossed 14 times to the C57Bl/6NCrl background prior to experimentation. *Cdk14^+/-^*mice were paired to generate *Cdk14^-/-^* mice. *PAC α-Syn*^A53T^ *TG* founder mice (dbl-PAC-Tg(*SNCA*^A53T^)^+/+^;*Snca*^-/-^, (28)) were provided by Robert L. Nussbaum (University of California, San Francisco, USA). Mice were genotyped using genomic DNA extracted from ear or tail tissue (genotyping protocol available upon request).

### Stereotactic PFF injections

Endotoxin-free recombinant mouse α-Syn fibril preparations were stored at −80 °C before usage (29,30). On the day of the stereotactic injections, fibrils were thawed and sonicated in a sonicator water bath to generate PFFs (Covaris S220, 18 W peak incident power, 20 % duty factor, 50 cycles per burst, 150 seconds). For transmission electron microscopy (TEM) analysis 0.2 mg/mL PFF samples were stained with Uranyl Acetate and imaged on an FEI Tecnai G2 Spirit Twin TEM (Centre for Advanced Materials Research [CAMaR], University of Ottawa). 6-month-old mice were deeply anesthetized with isoflurane, and 1 µL PFFs (5 mg/mL) or sterile saline (0.9 % NaCl) was unilaterally delivered into the dorsal striatum of the right hemisphere at these coordinates relative to bregma: −2 mm medial-lateral; +0.2 mm antero-posterior and −2.6 mm dorso-ventral. Injections were performed using a 2 µL syringe (Hamilton Company, Reno, NV, USA) at a rate of 0.1 µL/minute (min) with the needle left in place for at least 3 min before its slow withdrawal. After surgery, animals were monitored, and post-surgical care was provided. Behavioral experiments were performed 6 months post-injection, followed by the collection of the brains at 7 months post-injection.

### Intracerebroventricular administration of the CDK14 inhibitor FMF-04-159-2

Alzet® Mini-Osmotic Pumps (model 2004) were loaded with FMF-04-159-216 (R&D Systems 7158, Minneapolis, MN, USA) 1.47 μg/uL in vehicle solution containing 8 % DMSO (Fisher Scientific, BP231, Hampton, NH, USA), 2 % Tween 80 (Fisher Scientific, BP338-500) and 90 % ddH_2_O) 16 hours before stereotactic surgeries. Brain infusion catheters with 3.5 cm long catheter tubing (Alzet® Brain Infusion Kit) were attached as per manufacturer’s instructions. Brain infusion assemblies were incubated at 37 °C in sterile saline until implantation. 4-month-old *PAC α-Syn*^A53T^ *TG* were deeply anesthetized with isoflurane for the stereotactic implantation of brain infusion assemblies. FMF-04-159-2 (release rate of 0.35 mg/kg/day) or its vehicle solution was continuously administered into the cerebral ventricles (coordinates relative to bregma: −1.1 mm medial-lateral; −0.5 mm antero-posterior and −3 mm dorso-ventral) for 28 days with the brain infusion catheter attached to the skull and the connected pump in a subcutaneous pocket of the mouse’s back. The body weight of mice was measured and their activity, neurological signs, facial grimace, coat condition and respiration were scored (from 0 to 3) within 28 days after the surgery. Mice were sacrificed and organs were collected on the 28^th^ day of the administration period.

### Behavioral experiments

Grip strength tests were performed by holding the mice at an automatic grip strength meter (Chatillon DFE II, Columbus Instruments, Columbus, OH, USA) allowing them to grip the grid of the device with their fore- and hindlimbs. Then, mice were gently pulled back by their tail until they released the grip. Force exerted by the mouse during its removal from the grid, as measured by gram force (g), was evaluated 5 times per mouse. For nesting behavior tests, mice were singly caged overnight (16 hours) with a 5 cm x 5 cm cotton nestlet in a clean cage. Produced nests were scored on a scale from 1to 5 as previously described (31). For the tail suspension test, the tails of the mice were taped to a metal bar attached to the ENV-505TS Load Cell Amplifier and DIG-735 cabinet with high pass filter set to 1 Hz (Med Associates, Fairfax, VT, USA). The time of immobility was tested over 6 min. For the elevated plus maze test, mice were placed in the center of a maze consisting of two arms (6 cm x 75 cm), one open and the other enclosed. Over 10 min, the number of open arm entries was tracked with Ethovision software (Noldus Information Technology, Leesburg, VA, USA) and normalized to the total amount of arm entries. For the Y maze test, mice were placed in the center of the Y maze, where the three arms meet and given 8 min to explore. The number of arm alternations is measured relative to total arm entries using Ethovision software. For the open field test, mice were placed in a 45 cm x 45 cm open top cage and locomotion was automatically tracked with an overhead-mounted camera connected to a computer equipped with Ethovision tracking software (Noldus Information Technology). Open field motor activity was recorded over 10 min. For the hanging wire test, mice were placed on a wire cage lid which was gently turned upside down over a cage, followed by the recording of the latency to falling from the lid. Mice were given three consecutive training trials, followed by three test trials. The average latency to fall was normalized to the body weight of the mouse. Pole tests were performed by placing the mice on the top of a vertical pole (8 mm diameter and 55 cm height) with a rough surface. Mice were placed vertically, on the top of the pole, and the time required for turning was recorded. The mean time to turn was calculated from 5 consecutive trials for each mouse. For the rotarod test, mice were placed on a rotating, textured rod (IITC Life Science, Woodland Hills, CA, USA), with the speed gradually increasing from 4 to 40 rpm over 5 min. The latency to fall from the rotating rod was recorded for every mouse. Four trials per day with 10 min inter-trial intervals were performed for three consecutive days.

### Tissue harvesting and processing

For biochemical approaches mice were anesthetized with isoflurane (Fresenius Kabi, CP0406V2, Bad Homburg, Germany), and decapitated. Brain tissue of 5-month-old *PAC α-Syn*^A53T^ *TG* mice was weighed and lysed 1:3 (w/v) in PEPI buffer (5 mM EDTA, protease inhibitor [GenDEPOT, P3100, Katy, TX, USA] and phosphatase inhibitor [GenDEPOT, P3200] in PBS) with a Dounce homogenizer. Samples were further lysed 1:6 (w/v) using the tissue weight in TSS Buffer (140 mM NaCl, 5 mM Tris–HCl), then TXS Buffer (140 mM NaCl, 5 mM Tris–HCl, 0.5 % Triton X- 100), and SDS Buffer (140 mM NaCl, 5 mM Tris–HCl, 1 % SDS), as previously described (32). For immunohistology with paraffin sections, mice were anesthetized with 120 mg/kg Euthanyl (DIN00141704) and intracardially perfused with 10 mL of PBS, followed by 20 mL of 10 % Buffered Formalin Phosphate (Fisher Scientific, SF100-4). Brains were isolated and fixed in 10 % Buffered Formalin Phosphate at 4 °C for at least 24 hours. After dehydration by 70 %, 80 %, 90 % and 100 % ethanol and clearing by Xylenes, brains were infiltrated and embedded in paraffin (Louise Pelletier Histology Core Facility, University of Ottawa). Brains were sectioned at 5 µm.

### SDS-PAGE and mouse protein immunoblots

4X Laemmli buffer (Bio-Rad, 1610747, Hercules, CA, USA) with 20 % 2-mercaptoethanol (Bio-Rad, 1610710) was added to cleared protein and boiled at 95 °C for 5 min. Protein samples were loaded on a 12 % SDS-PAGE gel in the Mini-PROTEAN Tetra Cell (Bio-Rad, 165-8000). Protein was then transferred to a 0.2 µm nitrocellulose membrane (Bio-Rad, 1620112) using the Mini Trans-Blot Electrophoretic Transfer Cell (Bio-Rad, 1703930) in Tris-Glycine buffer with 10 % methanol (Fisher Scientific, A412P) at 340 mA for 90 min at 4 °C. Membranes were then blocked in 5 % milk in 1X TBS-T for 1 hour at room temperature followed by overnight incubation in primary antibody against pSer129 (pS) α-Syn (1:2 000, Abcam, 51253, Cambridge, UK), α-Syn (1:2 000, BD Biosciences, 610787, Franklin Lakes, NJ, USA), CDK14 (1:500, Santa Cruz Biotechnology, sc50475, Dallas, Texas, USA) and GAPDH (1:40 000, Proteintech, 60004-1-Ig, Rosemont, IL, USA) diluted in 2 % BSA, 0.02 % NaN_3_ in 1X TBS-T. Next, membranes were washed in TBS-T, followed by incubation in secondary antibody (peroxidase-conjugated donkey anti-rabbit IgG, (Cedarlane, 711-035-152, Burlington, ON, Canada) or donkey anti-mouse IgG, (Cedarlane, 715-035-150), both at 1:10 000 diluted in 5 % milk in TBS-T) for 1 hour at RT. Membranes were washed again in TBS-T, bathed in enhanced chemiluminescent reagent (Bio-Rad, 1705061), imaged using the ImageQuant LAS 4100 Imaging system (GE) and quantified using Image Lab 6.1 software (Bio-Rad).

### Dopamine Measurements with liquid chromatography-mass spectrometry/mass spectrometry (LC-MS/MS)

Striatal tissue punches from 3 mm thick brain sections from 12-month-old PFF-treated mice were weighed and submitted to The Metabolomics Innovation Centre (TMIC, Edmonton, AB, Canada) for analysis. Samples were homogenized in 50 µL of tissue extraction buffer, followed by centrifugation. Dopamine content in µM was analyzed by reverse-phase LC-MS/MS custom assay in combination with an ABSciex 4000 QTrap® tandem mass spectrometer (Applied Biosystems/ MDS Analytical Technologies, Foster City, CA, USA) using isotope-labeled internal standards. The assay utilizes a 96 deep well plate with a filter plate attached on top. Samples were thawed on ice, vortexed and centrifugated at 13 000 x g. 10 µL of sample was loaded in the center of the filter and dried in a stream of nitrogen. Following derivatization by phenyl isothiocyanate and drying of filter spots, dopamine content was extracted by adding 300 µL of extraction solvent, centrifugation into the lower collection plate and dilution by MS running solvent. Mass spectrometric analysis was performed with the ABSciex 4000 QTrap® tandem mass spectrometer in combination with an Agilent 1260 series UHPLC system (Agilent Technologies, Palo Alto, CA, USA). Samples were delivered by an LC method followed by a direct injection method. Data was analyzed using Analyst 1.6.2 and expressed as dopamine concentration relative to tissue weight.

### Histology

For Diaminobenzidine (DAB) antibody staining, paraffin sections were deparaffinized in xylenes and rehydrated in a series of decreasing ethanol (100 %, 90 %, 70 %, 50 %) followed by antigen retrieval in sodium citrate buffer (2.94 g sodium citrate, 0.5 mL Tween 20 in 1 L PBS, pH6) at 80 °C for 2 hours and quenching of endogenous peroxidase with 0.9 % H_2_O_2_ in PBS for 10 min. Sections were blocked in blocking buffer (0.1 % Triton X-100, 10 % normal horse serum in PBS) and incubated in primary antibody (pSer129 α-Syn, 1:500, Abcam, ab51253 or tyrosine hydroxylase, 1:500, Sigma-Aldrich, AB152, St. Louis, MO, USA) overnight at 4 °C. Then, sections were incubated in secondary antibody (donkey anti-rabbit biotin-conjugated, Jackson ImmunoResearch, 711-065-152, West Grove, PA, USA) 1:125 in blocking buffer for 2 hours and tertiary antibody solution (streptavidin-horseradish peroxidase conjugated, 1:250 in blocking buffer, Sigma-Aldrich, RPN1231V) for 2 hours before being exposed to DAB (Vector Laboratories, SK-4100, Newark, CA, USA) for 10 min. Hematoxylin counterstaining was conducted using the hematoxylin and eosin (H&E) Staining Kit (Abcam, ab245880) as per manufacturer’s instructions. Stained sections were dehydrated in a series of ethanol and xylenes solutions, followed by mounting sections with Permount (Fisher Scientific, SP15-100) and covering with coverslips.

H&E stainings were performed by the Louise Pelletier Histology Core Facility at the University of Ottawa with a Leica Autostainer XL (Leica Biosystems Inc, Concord, ON, Canada). After deparaffinization, sections were exposed to hematoxylin for 7 min and eosin for 30 sec. Then, sections were dehydrated, mounted and covered with coverslips.

For immunofluorescence antibody staining, tissue sections and primary neurons were incubated in primary antibody (Synapsin, 1:2 000, Thermo Fisher Scientific, Waltham, MA, USA, A-6442, PSD95, 1:500, Synaptic Systems, Göttingen, Germany, 124 011, pSer129 α-Syn, 1:500, Abcam, ab51253, Map2, 1:5 000, Abcam, ab5392) in blocking buffer overnight at 4 °C. Next, sections/neurons were incubated in secondary antibody (goat anti-rabbit IgG (H+L) Alexa Fluor^TM^ 647, Thermo Fisher Scientific, A-21244, for Synapsin, goat anti-mouse IgG (H+L) Alexa Fluor^TM^ 488, Thermo Fisher Scientific, A-11001, for PSD95, goat anti-rabbit IgG (H+L) Alexa Fluor^TM^ 488, Thermo Fisher Scientific, A-11008, for pS129 α-Syn and goat anti-chicken IgY (H+L) Alexa Fluor^TM^ 647, Thermo Fisher Scientific, A-21449, for Map2, all at 1:500) together with DAPI (Sigma-Aldrich, D9542) for 1 hour at RT, followed by mounting sections with fluorescence mounting medium (Agilent, S302380-2). Brightfield and epifluorescence micrographs were acquired using an Axio Scan Z1 Slide Scanner (Carl Zeiss AG, Oberkochen, Baden-Württemberg, Germany) (20x objective, Louise Pelletier Histology Core Facility) and a Zeiss AxioImager M2 (10x objective, Cell Biology and Image Acquisition Core Facility, University of Ottawa) and analyzed using ImageJ (National Institute of Health, Bathesda, MD, USA 1.52p) with 2-3 sections per mouse by a blinded investigator. Heatmaps were generated by counting pS129 α-Syn positive cells in defined regions of the brain (http://atlas.brain-map.org), normalized to the area occupied by the region and resulting cell densities were expressed as hues of red. Micrographs of *in situ* hybridization experiments for the expression of *Snca* (experiment 79908848) and *Cdk14* (experiment 71670684) in the mouse brain were downloaded from the Allen Brain Atlas (http://mouse.brain-map.org/) on August 3^rd^, 2022.

### Cell culture

Cells were kept at 37 °C, 5 % O_2_, 10 % CO_2_. For primary mouse cortical neurons coverglass #1.5 (Electron Microscopy Sciences, Hatfield, PA) were coated for at least 16 hours at 37 °C, washed twice with ddH_2_O and air dried at RT. Mouse embryos at embryonic days 14 to 16 were collected from pregnant females from *Cdk14*^+/-^ crossings (anesthetized with 120 mg/kg Euthanyl). Embryos were decapitated, brains removed, cortices isolated in Hank’s Balanced Salt Solution (HBSS) (Millipore Sigma, H9394-500ML) and dissociated in 0.7mg/ml trypsin (Sigma-Aldrich, T4549). Cell suspensions were quickly washed with 150 µg/mL trypsin inhibitor (Roche, Basel, Switzerland, 10109878001) and 666.67 µg/mL DNase 1 (Sigma-Aldrich, DN25-10mg), followed by another wash with 10 µg/mL trypsin inhibitor and 83.34 µg/mL DNase 1 and resuspended in neuronal media (Neurobasal Media (Thermo Fisher Scientific, Gibco 21103049), 1X B27 supplement (Gibco, 17504044), 1X N2 supplement (Thermo Fisher Scientific, Gibco 17502-048), 1X Penicillin Streptomycin (Thermo Fisher Scientific, Gibco SV30010), and 0.5 mM L-glutamine (Wisent Bio Products, Saint-Jean-Baptiste, QC, Canada, 609-065-EL). Single cells were seeded at ∼10^5^ cells per coverslip and maintained in culture for 9 (**Fig. S1D** and **S2B**) or 21 days (**Fig. 2**). Neurons were treated with 2 µg/mL of sonicated α-Syn PFFs (StressMarq, Victoria, BC, Canada, SPR-324) at 2 (**Fig. S1D** and **S2B**) or 7 days *in vitro* (DIV) (**Fig. 2**). 14 days after adding PFFs to the neuronal media, media was extracted and passed through a 40 µm cell strainer (VWR, Radnor, PA, USA, 21008-949) for *in vitro* α-Syn spreading experiments (**Fig. 2**). This conditioned media was added to untreated, naïve wildtype (WT) cortical neuron cultures at the time of the media change (7 DIV) and collected for immunofluorescence experiments at 21 DIV. Neurons treated with conditioned media were analyzed by immunofluorescence quantification after 14 days of treatment at 21 DIV. For immunofluorescence staining, cells were fixed in 4 % PFA for 20 min, washed in PBS and incubated in blocking buffer for 1 hour. Cells were incubated with primary antibodies overnight at 4 °C, followed by incubation with secondary antibodies for 1 hour at RT.

### hiPSC culture, neuronal differentiation, and Cas9-mediated gene editing

Human induced pluripotent stem cell (hiPSC) isogenic lines (Female) and human embryonic stem cell (hESC) isogenic lines (Male) were generated as described previously (33). hiPSCs were cultured as previously described (34) with slight modifications. Briefly, pluripotent cells were plated in mTeSR (Stem Cell Technologies, Vancouver, BC, Canada) and media was changed daily. The colonies were manually passaged weekly. Differentiation of hPSCs into A9-type DaNs was performed by following a floor plate differentiation paradigm (34,35). Immediately preceding differentiation, the colonies were dissociated into a single cell suspension using HyQTase. hPSCs were collected and re-plated at 4×10^4^ cells/cm^2^ on Matrigel (BD Biosciences)-coated tissue culture dishes for differentiation. Floor-plate induction was carried out using hESC-medium containing knockout serum replacement (KSR), LDN193189 (100 nM), SB431542 (10 μM), Sonic Hedgehog (SHH) C25II (100 ng/mL, Purmorphamine (2 μM), Fibroblast growth factor 8 (FGF8; 100 ng/mL), and CHIR99021 (3 μM). On day 5 of differentiation, KSR medium was incrementally shifted to N2 medium (25 %, 50 %, 75 %) every 2 days. On day 11, the medium was changed to Neurobasal/B27/Glutamax supplemented with CHIR. On day 13, CHIR was replaced with Brain Derived Neurotrophic Factor (BDNF; 20 ng/mL), ascorbic acid (0.2 mM), Glial Derived Neurotrophic Factor (GDNF; 20 ng/mL), transforming growth factor beta 3 (TGFβ3; 1 ng/mL), dibutyryl cAMP (dbcAMP; 0.5 mM), and DAPT (10 μM) for 9 days. On day 20, cells were dissociated using HyQTase and re-plated under high cell density 4×10^5^ cells/cm^2^ in terminal differentiation medium (NB/B27 + BDNF, ascorbic acid, GDNF, dbcAMP, TGFβ3 and DAPT) also referred to as DA Neuron (DAN)-Medium, on dishes pre-coated with poly-ornithine (15 μg/mL)/laminin (1 μg/mL)/fibronectin (2 μg/mL). Cells were differentiated for up to 60 DIV, with analysis being performed at DIV14, DIV 45 and/or DIV 60. At D10D and/or D14D of differentiation, hiPSC cultures were transduced with lentivirus containing Cas9 (lentiCRISPR v2, addgene, plasmid #52961) with the following gRNAs: non-targeting (5’- CGCTTCCGCGGCCCGTTCAA-3’), CDK14 exon 3 (5’-GCAAAGAGTCACCTAAAGTT-3’) and exon 8 (5’-TGTGCAAAATATAACGCTGG-3’). From D10D-D14D media was supplemented with 0.1 µM compound E (AlfaAesar, J65131, Haverhill, MA, USA). At D18D, cells were replated onto poly-ornithine (15 μg/mL)/laminin (1 μg/mL)/fibronectin (2 μg/mL) coated plates. Cells were maintained in DAN-medium (DMEM/F12, 200uM Ascorbic Acid, 0.5 mM dbcAMP, 20 ng/mL BDNF, 20 ng/mL GDNF, 1 ng/mL TGFb3, and 1 % Anti-Anti) for 6-7 weeks where lysates were then collected for protein analysis.

### hESC culture, neuronal differentiation and *in vitro* CDK14 inhibitor treatment

Neural progenitor cell (NPC) differentiation was performed as previously described (36). Dishes for hESC cultures were coated with 0.15 mg/mL growth factor reduced Matrigel (Corning Inc, 354230, Corning, NY, USA) in DMEM/F-12 (Thermo Fisher Scientific, 11320033) for 1 hour at RT prior to cell seeding. H9 hESCs (WiCell, WA09, Madison, WI, USA) were seeded as colonies and maintained in mTeSR Plus (StemCell Technologies, 05825). NPC differentiation was initiated when the hESC cultures reached 90 % confluence, by replacing growth medium with Knockout Serum Replacement (KSR) medium (414 mL Knockout-DMEM (Thermo Fisher Scientific, 10829018), 75 mL Knockout-serum replacement (Thermo Fisher Scientific, 10828028), 5 mL Glutamax (Thermo Fisher Scientific, 35050061), 5 mL MEM-NEAA (Thermo Fisher Scientific, 11140050), 500 µL 2-mercaptoethanol (Thermo Fisher Scientific, 21985023), 500 µL Gentamicin (Wisent Bioproducts 450-135), 10 µM SB431542 (Tocris, 1614, Bristol, UK) and 500 nM LDN- 193189 (Stemgent 04-0074, Reprocell Inc, Yokohama, Kanagawa 222-0033, Japan). Differentiation medium was replaced daily on days 4 and 5 by 75:25 KSR:N2 medium (486.5 mL DMEM/F-12 (Thermo Fisher Scientific, 11320033), 5 mL 15 % glucose, 5 mL N2 supplement (Thermo Fisher Scientific, 17502048), 500 µL 20 mg/mL human insulin (Wisent Bioproducts, 511-016-CM), 2.5 mL 1M HEPES (Thermo Fisher Scientific, 15630080), 500 µL Gentamicin), on days 6 and 7 by 50:50 KSR:N2, on days 8 and 9 by 25:75 KSR:N2 and on days 10 and 11 by N2 medium containing 500 nM LDN-193189. On day 12, differentiated NPCs were treated with Y-27632 (Tocris, 1254) for 4 hours, dissociated with Accutase (Stemcell Technologies, 07922) and seeded into Matrigel coated dishes containing Neural Induction Medium (NIM, 244 mL DMEM/F12, 244 mL Neurobasal medium (Thermo Fisher Scientific, 21103049), 2.5 mL N2 Supplement, 5 mL B-27 Supplement (Thermo Fisher Scientific, 17504044), 2.5 mL GlutaMAX^TM^ (Thermo Fisher Scientific, 35050061), 125 µL 20 mg/mL human insulin, 500 µL 20 µg/mL FGF2 (StemBeads, SB500, Rensselaer, NY, USA), 10 µL 1 mg/mL hEGF (Millipore Sigma E9644) and 500 µL Gentamicin) for expansion. NPCs were passaged at full confluence a minimum of one time before neuronal differentiation.

For NPC-neuronal differentiation culture dishes were coated with 0.001 % Poly-L-ornithine (Millipore Sigma, P4957) at 4 °C overnight, followed by 25 µg/mL laminin (Millipore Sigma, L2020) for 2 hours at room temperature. NPCs were treated with Y-27632 for 4 hours, dissociated with Accutase and seeded at a density of 20 000 cells/cm^2^ in NIM. Neuronal differentiation was initiated when NPCs reached 70 % confluence by replacing growth medium with neuronal differentiation medium (244 mL DMEM/F-12 medium, 244 mL Neurobasal medium, 2.5 mL N2 supplement, 5 mL B27 supplement, 200 µL 50 µg/ml BDNF (Peprotech, 450-02), 200 µL 50 µg/ml GDNF (Peprotech 450-10, Thermo Fisher Scientific), 250 mg dibutyryl cyclic-AMP (Millipore Sigma, D0627), 500 µL 100 M L-ascorbic acid (FujiFim Wako Chemicals, 323-44822, Osaka, Japan), and 500 µL Gentamicin). Cells were fed every 3 days for 18 days to obtain immature neuronal networks. FMF-04-159-2 was dissolved in DMSO (Fisher Scientific, BP231) and applied in cell culture medium to hESC-derived neurons for 6 days (with a replenishment of FMF-04-159- 2 -containing medium after the first 3 days). For protein analysis cells were washed with cold PBS, scraped, and collected in low protein binding microcentrifuge tubes (Thermo Scientific, 90410). Cells were pelleted by centrifugation at 1 000 x g for 5 min at 4 °C. The supernatant was aspirated, and the cells were lysed in cold RIPA buffer (50 mM Tris, pH 7.5, 150 mM NaCl, 0.1 % SDS, 0.5 % sodium deoxycholate; 1 % NP-40, 5 mM EDTA, pH 8.0) with protease and phosphatase inhibitors. Cell lysates were incubated on ice for 20 min, with vortexing every 5 min. Lysates were centrifuged at 18 000 x *g* for 20 min at 4 °C to pellet cell debris.

### ELISA (Enzyme-Linked ImmunoSorbent Assay)

ELISA α-Syn protein quantification was performed as previously described (37,38). 384-well MaxiSorp plates (Nunc, Inc) were coated with capturing antibody (α-Syn, BD Biosciences, 610787) diluted 1:500 in coating buffer (NaHCO_3_ with 0.2 % NaN3, pH9.6) overnight at 4 °C. Following 3 washes with PBS/0.05 % Tween 20 (PBS-T), plates were blocked for 1 hour at 37 °C in blocking buffer (1.125 % fish skin gelatin; PBS-T). After 3 washes, samples were loaded in duplicates and incubated at RT for 2 hours. Biotinylated hSA4 antibody (in-house antibody) was generated using 200 μg Sulfo-NHS-LC Biotin (Pierce, Thermo Fisher Scientific), diluted 1:200 in blocking buffer and added to the plate for 1 hour at 37 °C. Following 5 washes, ExtrAvidin phosphatase (Sigma, E2636) diluted in blocking buffer was applied for 30 min at 37 °C. Color development was carried out by using fast-p-nitrophenyl phosphate (Sigma, N1891) and monitored at 405 nm every 2.5 min for up to 60 min. Saturation kinetics were examined for identification of time point(s) where standards and sample dilutions were in the log phase.

### Primary rat neurons, human α-Syn PFFs and protein analysis

Cortical neurons were harvested from the E18 Sprague Dawley rat embryos (Charles River, Wilmington, MA, USA). The harvested cortical tissue was digested using 17 U/mg Papain followed by mechanical dissociated by gentle trituration through a glass flamed Pasteur pipet. The cells were seeded into plates coated 24 hours prior to dissection with Poly-D-Lysine (0.15 mg/mL). The cells were incubated at 37 °C, 7.5 % CO_2_ until collection. Every 3 to 4 days, a 50 % media change was performed (2 % B27 supplement, 1 % antibiotic/antimycotic, 0.7 % BSA Fraction V, 0.1 % β-mercaptoethanol in HEPES-buffered DMEM/F12). Where required, cells were exposed to 100 nM FMF-04-159-2 (Bio-Techne, 7158/10, Minneapolis, MN, USA) dissolved in DMSO, at 14 DIV, and again at the subsequent feed (18 DIV). At 14 DIV, cells were exposed to either 1 μg/mL human α-Syn PFFs, or 1 μg/mL monomeric α-Syn. Cell lysates were collected at day 5 post PFF or monomeric exposure. Human α-Syn protein was isolated from BL21-CodonPlus (DE3)-RIPL competent cells transformed with pET21a-alpha-synuclein and purified by Reversed-phase HPLC. PFFs were then generated as previously described (39). Purified α-Syn (5 μg/mL in PBS) was incubated at 37 °C with constant shaking for 7 days, then aliquot and stored at −80 °C. Prior to use, PFFs were thawed and diluted in PBS, then subjected to sonication (20% amplitude, 30 seconds; 1 second on, 1 second off) and added to neuronal media for exposure to neurons at a concentration of 1 μg/mL for 24 hours. Following the incubation, cell lysates were collected in 150 μL ice-cold RIPA buffer containing phosphatase and protease inhibitors (1 mM aprotinin, 1 mM sodium orthovanadate, 1 nM sodium fluoride, and 10 mM phenylmethylsulfonyl fluoride). Samples were homogenized using an 18G needle, left on ice to rest for 15 min, and then centrifuged at 14 000 *g* to remove any cellular debris. For the soluble fraction, cells were lysed in 1 % Tx-100 in TBS buffer (TXS buffer) and cleared by ultracentrifugation at 100 000g for 30 min. Pellets were washed twice with 1 % Tx-100 in TBS, then resuspended in 8M Urea + 8 % SDS in TBS buffer (Urea buffer) to generate the insoluble fraction. Using the BioRad DC Protein Assay kit, the protein concentration of each sample was quantified following the manufacturer’s guidelines. SDS-PAGE was performed using 12.5 % resolving gels and 4 % stacking gels, and gels were run for 15 min at 80 V followed by approximately 1.5 hours at 110 V. The gels were transferred onto 0.2 μM nitrocellulose membranes at 35 V and 4 °C overnight. Following the transfer, the membranes were blocked for 1 hour at room temperature using blocking buffer (5 % non-fat dry milk in 1 X TBST) with constant agitation. Primary antibodies were prepared in blocking buffer containing 0.1 % Tween 20 and were probed overnight at 4 °C under constant agitation (CDK14, 1:1 000, Santa Cruz Biotechnology, sc50475; TH, 1:1 000, Pel Freeze Biologicals, Rogers, AR, USA, P40101; a-Syn, 1:1 000, BD Biosciences, 610787; pS129 a-Syn, 1:500, abcam, ab51253; β-Actin, 1:1 000, rabbit, Biolegend, San Diego, CA, USA, 622101 or mouse, Sigma, A5411, βIII-Tubulin, 1:5 000, rabbit, Biolegend, 802001). Following primary antibody incubation, membranes were rinsed using 1XPBS containing 0.1 % Tween 20 and subsequently re-blocked using the blocking buffer. The membranes were then probed with secondary antibody for 1 hour at RT in blocking buffer containing 0.1 % Tween 20 (Goat anti-Mouse IgG (H+L) Secondary Antibody, HRP (Thermo Fisher Scientific, 31430); Goat anti-Rabbit IgG (H+L) Secondary Antibody, HRP (Thermo Fisher Scientific, 31460); Li-Cor infrared conjugated secondary/ IRDye 800RD Donkey anti-Rabbit IgG antibody (LI-COR Biosciences, 926-32211, Lincoln, NE, USA) at dilutions of 1:2 000). The membranes were rinsed to remove any residual blocking buffer using 1X PBS containing 0.1 % Tween 20. If HRP-conjugated secondary antibodies were used, membranes were probed for 5 minutes with clarity Western enhanced chemiluminescence blotting substrate (Bio-Rad) and visualized with photosensitive film. For LiCOR-secondary antibodies, membranes were visualized with a LiCOR Odyssey Fc. Total protein was visualized by Coomassie Brilliant Blue (0.1 % Coomassie, 50 % Methanol and 10 % glacial acetic acid in ddH_2_O).

### Statistics

Statistical analysis was performed using GraphPad Prism version 9.2.0. Quantified data are visualized as mean + standard error of the mean (SEM). Unpaired student’s *t* tests were used for two-group comparisons. Data affected by one or two factors were analyzed by one-way or two-way analysis of variance (ANOVA), respectively, followed by Bonferroni *post hoc* comparisons (unless otherwise stated in the figure legend) when at least one of the main factors or the interaction was significant. A significance level of 0.05 was accepted for all tests. Asterisks mark *P* values ≤ 0.05 (*), ≤ 0.01 (**), ≤ 0.001 (***), or ≤ 0.001 (****).

## RESULTS

### CDK14 ablation limits grip strength deficits and reduces cortical α-Syn pathology in PFF-injected mice

We first examined existing *in situ* hybridization data for the expression of *Snca* and *Cdk14* in the murine brain. We observed that both genes are expressed in similar brain regions, including the hippocampus and the SN (**Fig. S1A**). Since PD is a chronic disease, inhibition of a candidate modifier would have to be safe in the long term. We analyzed Cdk14 protein levels in different mouse organs and tested the effects of Cdk14 depletion on survival, fertility, and organ cytoarchitecture *in vivo*. We found that CDK14 protein is highly abundant in the brain, as well as in the lung and the spleen (**Fig. S1B**). Cdk14 nullizygous mice are viable, fertile, and exhibited normal brain morphology (**Fig. S1C**, (27)) and synaptic integrity (**Fig. S1D**). Furthermore, we did not observe altered morphology of the lung and spleen by Cdk14 ablation (**Fig. S1E**). We next asked whether silencing *Cdk14* is sufficient to mitigate behavioral and histological phenotypes observed in cultured neurons and in mice exposed to pathogenic α-Syn pre-formed fibrils (mouse PFFs; **Fig. 1A** and **B**; **Fig. S2A**). 6 months following intrastriatal injection of α-Syn PFFs, there is a stereotypic brain-wide accumulation of pS129 α-Syn – a marker of human synucleinopathies – in addition to SN DaNs loss and mild motor impairments (40–42). In our experimental paradigm, 6 months after PFF injection (at an age of 12 months), we found that PFF injected WT mice exhibited reduced forelimb force generation in the grip strength test compared to their saline-treated counterparts (**Fig. 1B**), similar to what has been previously reported (41,42). In contrast, the PFF-induced weakening of grip strength was not observed in *Cdk14*^+/-^ or in *Cdk14*^-/-^ mice. We did not observe PFF-mediated changes (in any genotype tested) in the other 8 behavioral tests conducted (including tests for cognitive and motor function). Importantly, we noted that saline-injected *Cdk14*^+/-^ and *Cdk14*^-/-^ mice consistently performed like their WT counterparts in each test, suggesting that chronic Cdk14 reduction is not deleterious to the brain (**Fig. S2C**).

**Figure 1.**
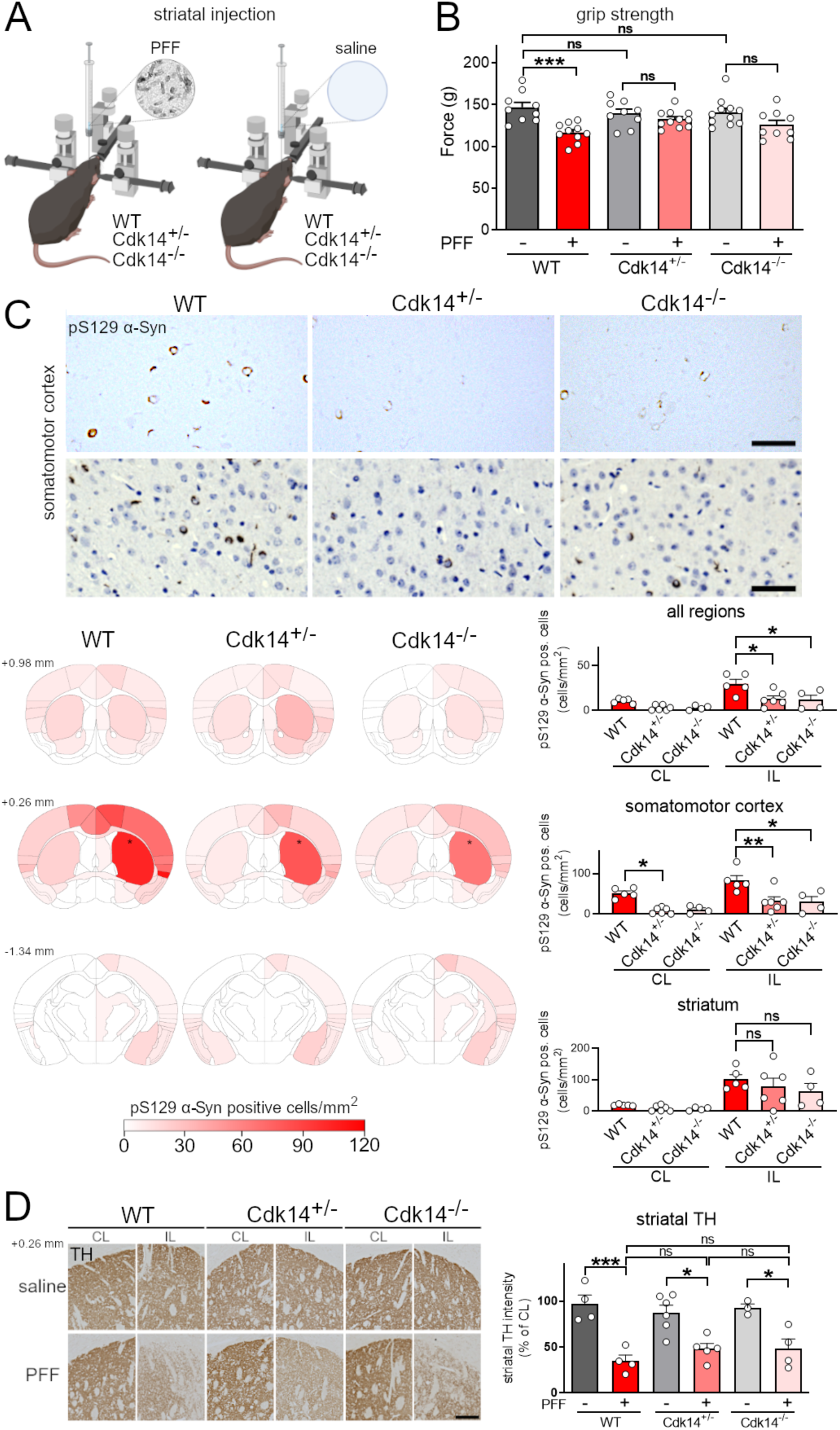
Loss of Cdk14 ameliorates grip strength impairment and α-Syn pathology in PFF- injected mice. (A) Mouse α-Syn PFFs were injected unilaterally into the striatum by stereotactic injection in 6-month-old mice. (B) Loss of forelimb grip strength at 6 months post α-Syn PFF-treatment in WT mice, but not in *Cdk14*^+/-^ or *Cdk14*^-/-^ mice. Mean + SEM, two-way ANOVA, Bonferroni *post hoc,* n: 9-10. (C) α-Syn PFF-injection increased the load of pS129 α-Syn-positive cells (depicted without and with hematoxylin counterstaining, 50 µm scale bars) in the injected hemisphere (ipsilateral, IL; injection site indicated by *). Densities of pS129 α-Syn-positive cells are represented in heat maps as hues of red at 3 rostrocaudal levels (relative to bregma: +0.98 mm, +0.26 mm, and −1.34 mm) with dark shades of red correlating to high cell densities (injection sites marked by asterisks). Amounts of pS129 α-Syn-positive cells in all brain regions (averaged in IL and contralateral to the injection, CL), the somatomotor cortex and the striatum at +0.26 mm relative to bregma of PFF-injected *Cdk14^+/-^* and *Cdk14^-/-^* mice were lower relative to their wildtype (WT) counterparts. Mean + SEM, two-way ANOVA, Bonferroni *post hoc* comparisons, n: 3-6. (D) PFF injection reduces the tyrosine hydroxylase (TH)-positive fiber density IL in comparison to the non-injected hemisphere (CL) to a similar degree in WT, *Cdk14^+/-^* and *Cdk14^-/-^* mice at +0.26 mm relative to bregma (200 µm scale bar). Mean + SEM, two-way ANOVA, Bonferroni *post hoc* comparisons, n: 3-7.

Next, we analyzed the relative pathological burden of accumulated α-Syn throughout the brain of mice injected with PFFs. We stained for synucleinopathy-linked pS129 α-Syn in the brain of WT mice and found high amounts of pS129 α-Syn-positive cells in the PFF-injected hemisphere (ipsilateral to the injection, IL), which were absent in saline-injected controls (data not shown). We then mapped the distribution of α-Syn pathology at three different rostrocaudal levels near the injection site (relative to bregma: +0.98 mm, +0.26 mm and −1.34 mm) and found an overall blunting of pS129 α-Syn-positive pathology in the PFF-injected *Cdk14*^+/-^ and *Cdk14*^-/-^ mice compared to their WT littermates (**Fig. 1C**). This was particularly evident at anatomical regions distal from the injection site, such as the somatomotor cortex (bregma +0.26 mm) and in other cortical areas. Surprisingly, α-Syn pathology proximal to the injection site in the striatum was not significantly affected between PFF-injected genotypes (**Fig. 1C**). Similarly to cortical regions of PFF-injected *Cdk14*^-/-^ mice, we observed fewer pS129 α-Syn-positive neurons in *Cdk14*^-/-^ mouse primary cortical cultures treated with PFFs compared to their WT littermate controls (**Fig. S2B**).

Previous reports have shown that α-Syn PFFs can induce nigrostriatal degeneration over time (40–42). We stained for tyrosine hydroxylase (TH) at the injection site in the striatum and observed a similar decrease of TH-positive dopamine fibers in WT, *Cdk14*^+/-^ and *Cdk14*^-/-^ mice injected with PFFs ipsilateral to the injection, relative to their saline-treated counterparts (**Fig. 1D**). PFF-induced loss of dopamine fibers was not accompanied by altered striatal dopamine content (**Fig. S2D**). TH staining in the midbrain revealed equal loss of SN DaNs in PFF-injected WT, *Cdk14*^+/-^ and *Cdk14*^-/-^ mice in comparison to saline-injected controls (**Fig. S2D**). Together, these data show that loss of Cdk14 blunts cortical α-Syn histopathology and PFF-induced grip strength impairment without evidently halting nigrostriatal neurodegeneration in the PFF model. It was interesting to note that the effect of Cdk14 on α-Syn was selective to pathological forms of the protein, as partial reduction or ablation of Cdk14 in *Cdk14^+/-^* or *Cdk14^-/-^* mice, respectively, did not change levels of endogenous mouse α-Syn (**Fig. S2E**).

### Cdk14 loss decreases α-Syn cell-to-cell spread in cortical neuron culture

Our *in vivo* data suggest that decreasing Cdk14 reduces distal (e.g. contralateral; higher cortical area) pathology, while not affecting pathology close to the injection site. We hypothesized that Cdk14 may preferentially affect the cell-to-cell spread of α-Syn, thus contributing to this phenomenon. To test this, we cultured primary neurons of WT, *Cdk14*^+/-^ and *Cdk14*^-/-^ embryos, treated them with mouse PFFs at 7 days in culture and collected media 14 days post PFF application (a time where most of the exogenous fibrils have been depleted from the system ((43); schematic in **Fig. 2A**). We applied this filtered, seed-competent media to naïve WT cultures to test the seeding capacity of α-Syn and found that loss of Cdk14 dramatically reduced α-Syn pathology (measured by pS129 α-Syn accumulation), 14 days following media application (**Fig. 2B**).

**Figure 2.**
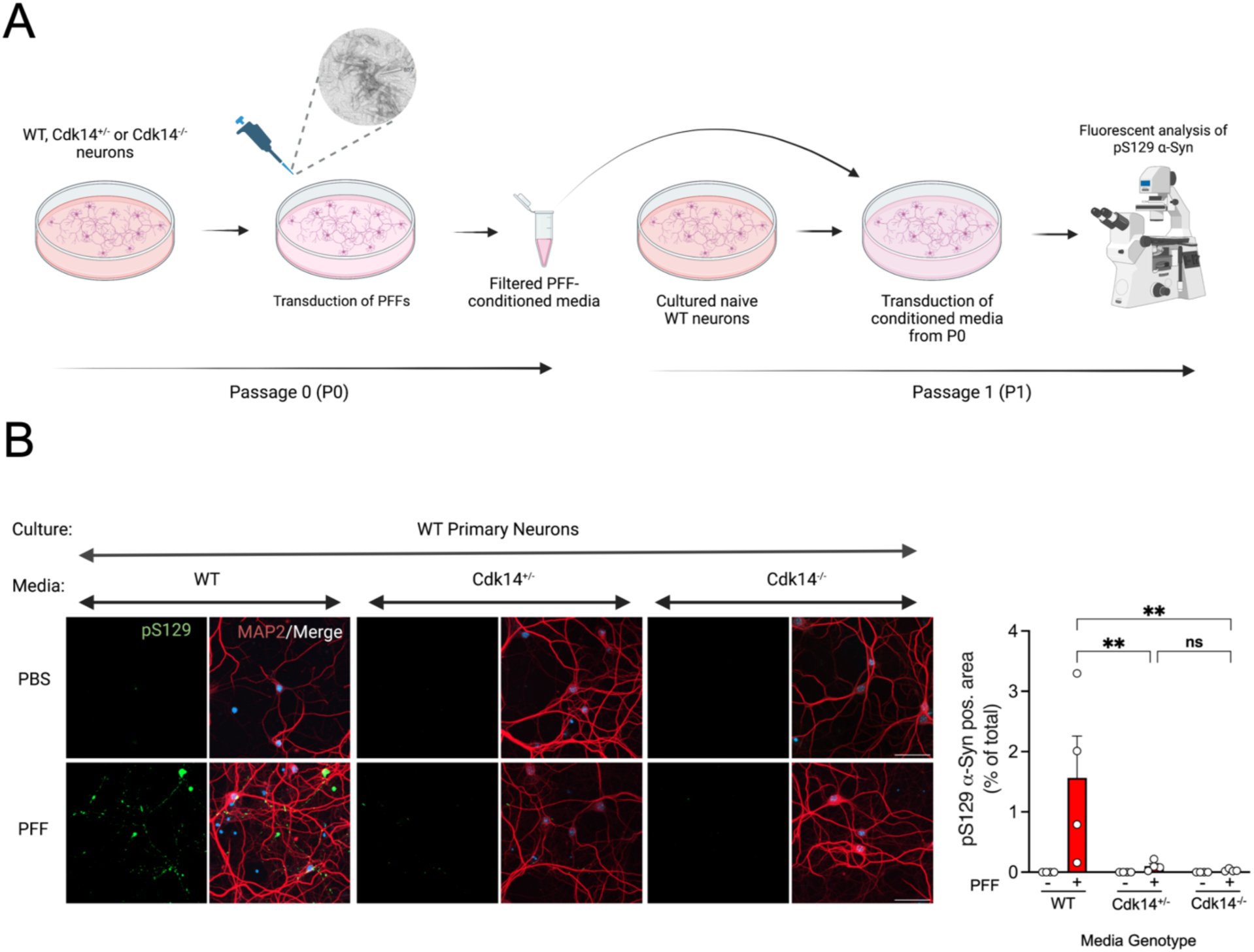
Primary cortical neurons lacking Cdk14 exhibit decreased spreading of seed-competent α-Syn. (A) Paradigm for passaging PFF-transduced WT, *Cdk14*^+/-^, and *Cdk14*^-/-^ primary culture media to WT, naïve cultures. Media from 21 DIV cultures transduced with PFFs for 14 days were collected, filtered, and added to naïve cultures at 7 DIV. Media-treated cultures were then fixed and analyzed for fluorescent pS129 α-Syn signal. (B) Conditioned media from *Cdk14*^+/-^ and *Cdk14*^-/-^ neurons induce less α-Syn pathology (indicated by pS129 α-Syn-positive area) in naïve WT neurons compared to WT conditioned media (50 µm scale bars). Mean + SEM, two-way ANOVA, Holm- Šídák post hoc comparisons, n=4.

### Knockdown of CDK14 decreases pS129 α-Syn levels in human neurons

Having observed the benefits of Cdk14 depletion in the PFF mouse model of PD, we next tested whether this benefit translates to human neurons. We infected DaNs derived from a PD patient carrying an A53T mutation in α-Syn (33) as well as its isogenic control with lentiviruses carrying Cas9/sgRNAs against *CDK14*. Neurons infected with sgRNAs targeting either exon 3 (E3) or exon 8 (E8) of *CDK14*, exhibited approximately 50 % of the CDK14 levels of the control cultures (**Fig. 3**). We found that A53T mutant cells show a marked increase of pS129 α-Syn compared to isogenic controls, and that *CDK14* knockdown significantly lowers pS129 α-Syn levels (**Fig. 3**).

**Figure 3.**
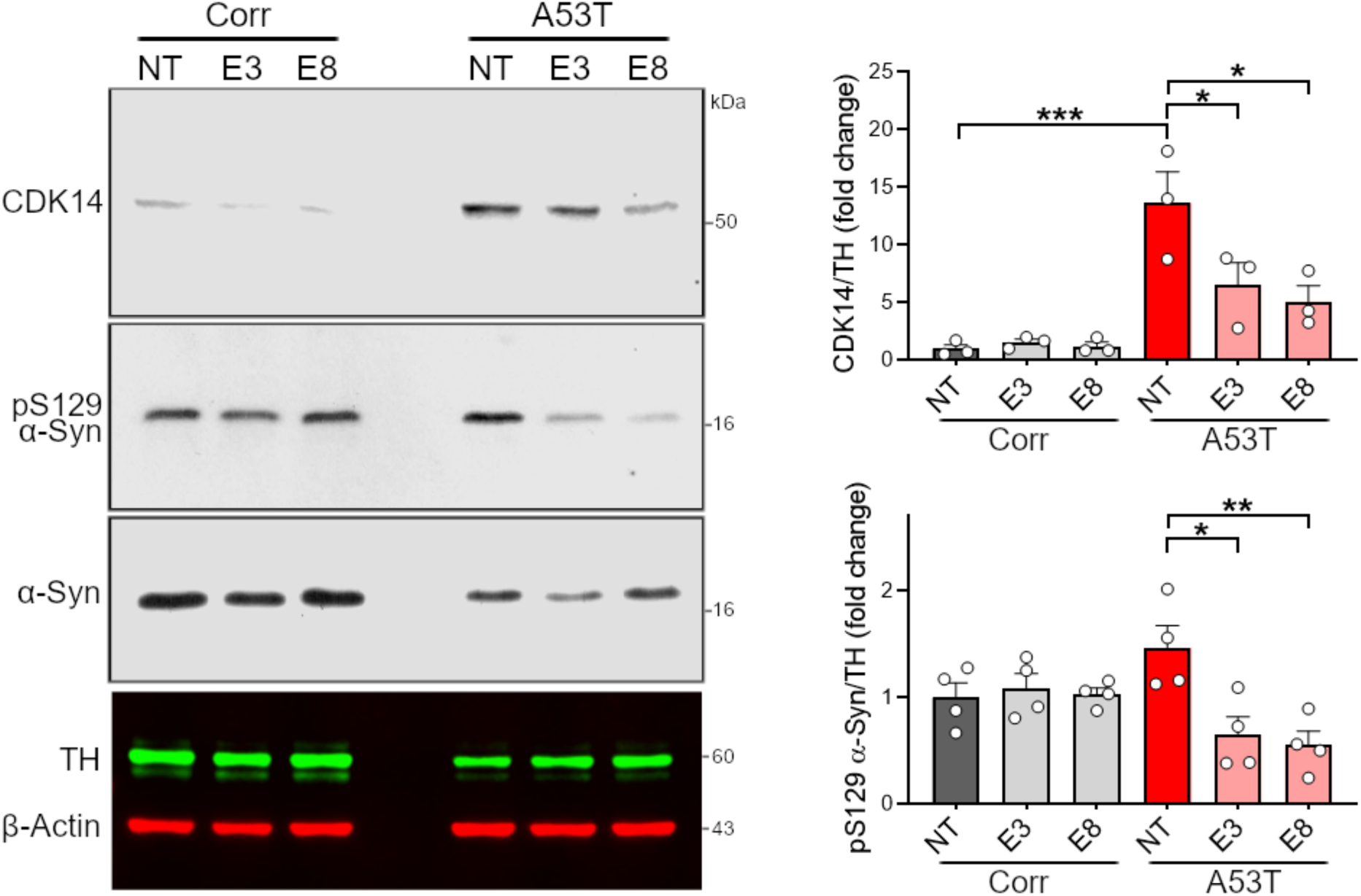
Genetic reduction of Cdk14 attenuates pS129 α-Syn in human neurons. Immunoblots illustrating elevated amounts of pS129 α-Syn in the RIPA buffer-soluble protein fraction of hiPSC-derived *SNCA* A53T human neurons (in comparison to isogenic corrected neurons [Corr]) which is reduced by CRISPR/Cas9-mediated knockdown of *CDK14* (targeted against *CDK14* exon 3 (E3) and 8 (E8), NT non-targeting control). Mean + SEM, one-way ANOVA, Tukey *post hoc* comparisons, n=3-4.

### Pharmacological targeting of CDK14 decreases α-Syn levels and mitigates its pathogenic accumulation

As kinases are typically druggable targets which can be inhibited in non-invasive ways (44), we next asked whether CDK14 inhibition would be a tractable route for decreasing α-Syn levels. We used a recently developed CDK14 covalent inhibitor (FMF-04-159-2) (26) to test whether acute inhibition of CDK14 reduces α-Syn levels. We treated hESC-derived cortical neurons with the CDK14 inhibitor for 6 days and observed a dose-dependent reduction in total α-Syn concentration by ELISA (**Fig. 4A**). We also tested whether the CDK14 inhibitor affects the α-Syn load in rat primary neuronal cultures treated with α-Syn PFFs. Here, PFF treatment induced a spike in insoluble (Urea buffer-soluble) α-Syn protein, which CDK14 inhibition markedly reduced (**Fig. 4B**). Interestingly, the PFF treatment also induced an increase of CDK14 insoluble protein, which was not present in untreated neurons or in neurons treated with α-Syn monomers (**Fig. 4B**).

**Figure 4.**
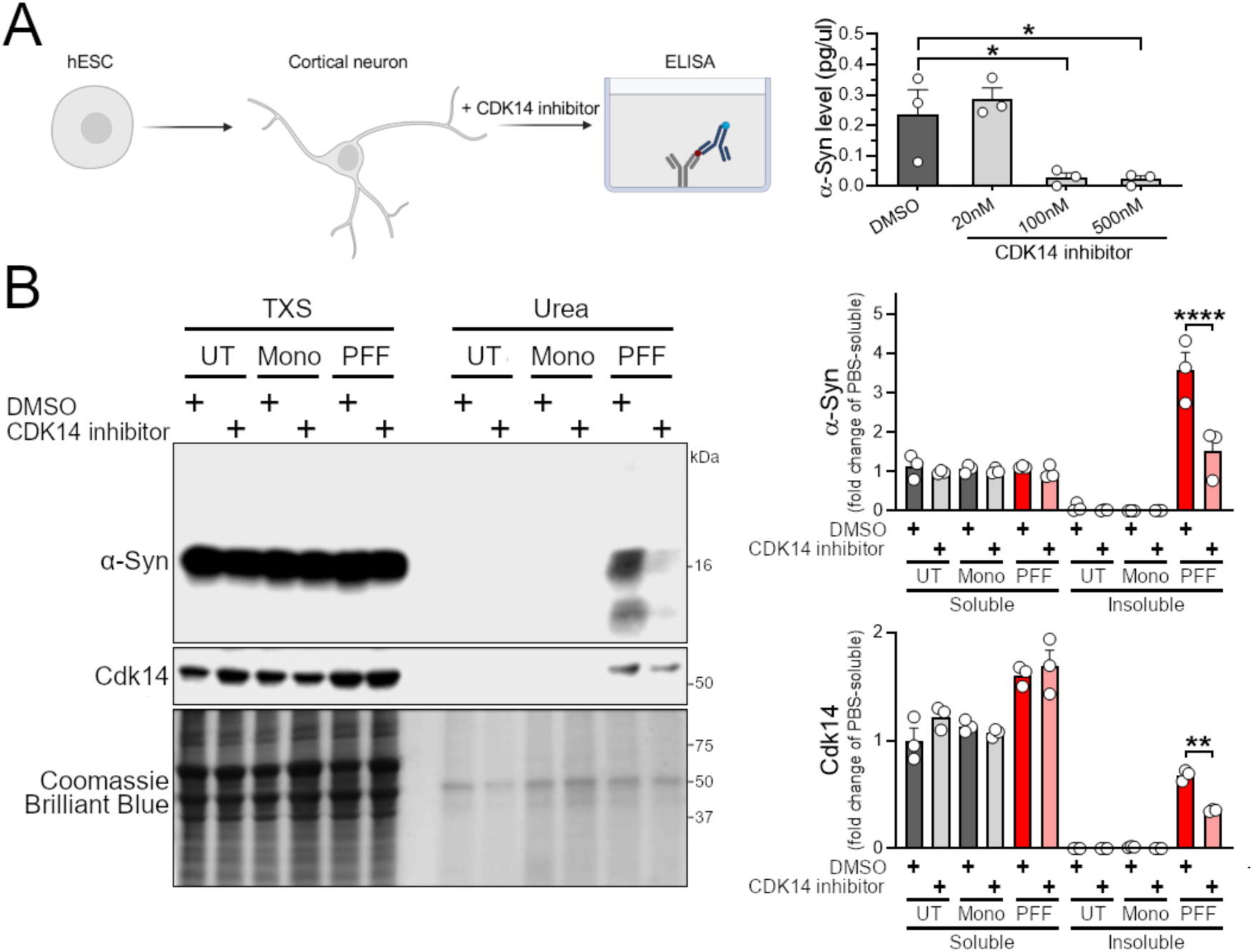
Pharmacological inhibition of CDK14 reduces α-Syn protein burden in human and rodent neurons. (A) Dose-dependent reduction of α-Syn in hESC-derived human neurons is detected by ELISA-quantification after 6 days of CDK14 inhibitor FMF-04-159-2 treatment. Mean + SEM, one-way ANOVA, Bonferroni *post hoc* comparisons, n=3. (B) Human α-Syn PFFs applied to rat cortical neurons for 5 days increase the amounts of α-Syn and CDK14 in the insoluble protein fraction (Urea buffer-soluble) compared to untreated (UT) and α-Syn monomers (Mono)-treated neurons, as shown by immunoblots, which were reduced by the application of 100 nM of the CDK14 inhibitor. Mean + SEM, two-way ANOVA, Holm-Šídák *post hoc* comparisons, n=3.

### *In vivo* inhibition of CDK14 decreases the load of human α-Syn

Since the treatment of human neurons and PFF-challenged rat neurons with the CDK14 inhibitor showed a reduction in total and insoluble α-Syn protein, respectively, we next tested whether pharmacological inhibition of CDK14 modifies α-Syn levels *in vivo*. We administered FMF-04- 159-2 via intracerebroventricular infusion at 0.35 mg/kg/day for 28 days in 4-month-old *PAC α-Syn*^A53T^ *TG* mice (**Fig. 5A**) which harbor the PD-associated A53T mutant human α-Syn gene in the absence of mouse *Snca* (28). Administration of the CDK14 inhibitor did not modify body weight development, nor induce any signs of distress or pain as indicated by alterations of locomotion, facial expression, or coat condition of *PAC α-Syn*^A53T^ *TG* mice in comparison to vehicle-treated counterparts (**Fig. S3A**). Similarly, CDK14 inhibitor treatment did not induce changes in the cytoarchitecture of the lung, spleen, and liver (**Fig. S3B**). To quantify levels of pathogenic forms α-Syn, we collected brains after 1 month of CDK14 inhibitor treatment (**Fig. 5A**) and analyzed protein content of the TSS, TXS and SDS buffer-soluble fractions (**Fig. 5B**). We observed a reduction of total α-Syn and low molecular weight α-Syn species, suggestive of decreased C-terminally truncated (CTT) α-Syn in CDK14 inhibitor-treated mice relative to their vehicle controls. Interestingly, levels of pS129 α-Syn were increased in CDK14 inhibitor treated mice in the TSS buffer-soluble fraction, whereas it was decreased in the TXS buffer-soluble fraction. Notably, CDK14 levels were specifically decreased (target engagement) in the TXS and SDS buffer-soluble fractions, but not the TSS buffer-soluble fraction, of inhibitor-treated mice. Together these results show that *in vivo* administration of the CDK14 inhibitor in the brain engages its target and mitigates certain pathogenic forms of human α-Syn in the mouse brain in a protein fraction-dependent manner without inducing obvious discomfort or pain.

**Figure 5.**
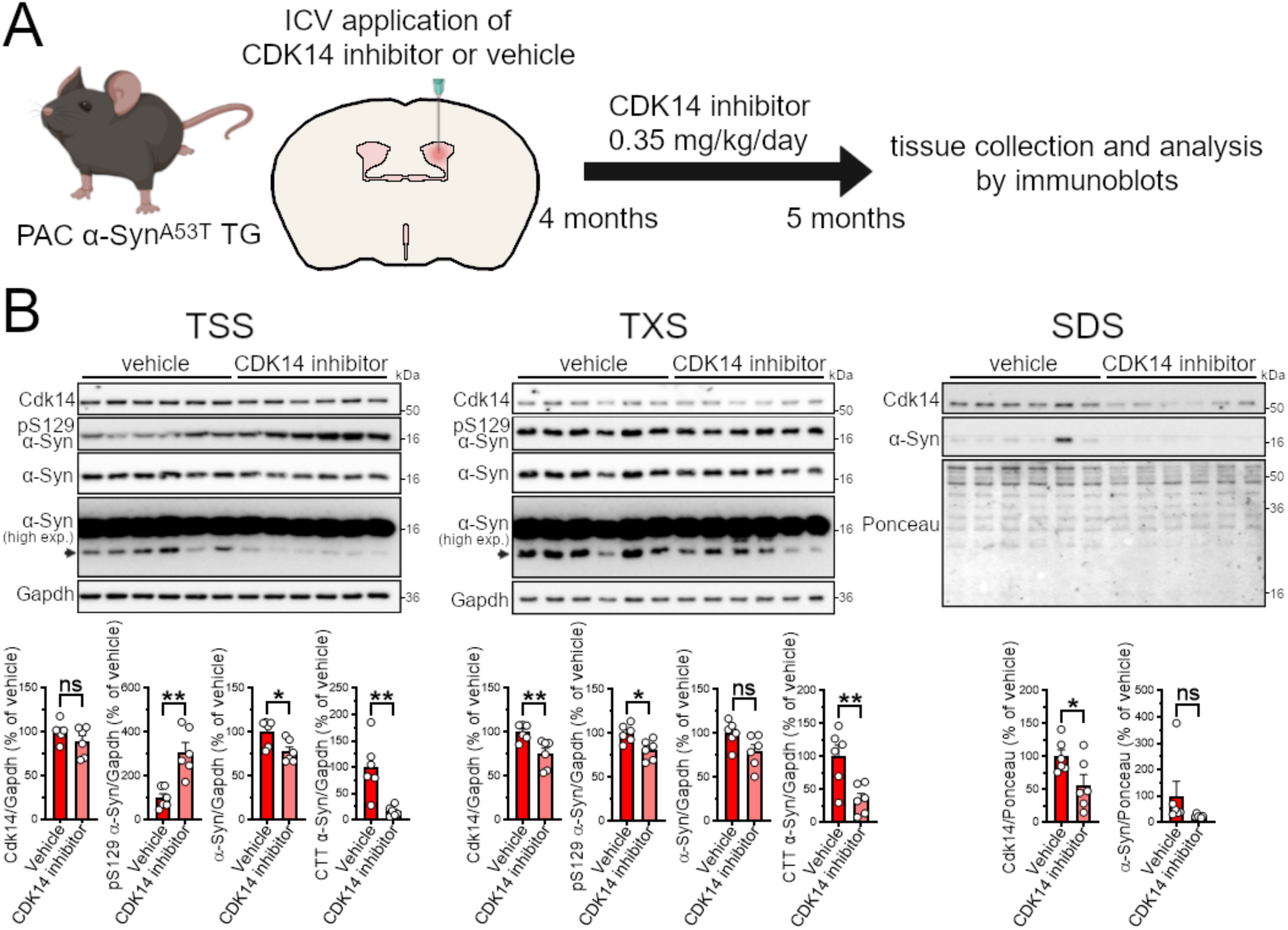
Inhibition of Cdk14 decreases human α-Syn protein in mice expressing A53T *SNCA*. (A) Intracerebroventricular (ICV) administration strategy for the CDK14 inhibitor FMF- 04-159-2 at 0.35 mg/kg/day for 28 days in 4-month-old *PAC α-Syn*^A53T^ *TG* mice. (B) Immunoblots with protein isolated from the brain of FMF-04-159-2-treated *PAC α-Syn*^A53T^ *TG* mice visualize the reduction of α-Syn in the TSS buffer-soluble, Cdk14 in the TXS buffer and SDS buffer-soluble and C-terminally truncated (CTT) α-Syn (arrowhead at high exposure (high exp.)) in the TSS buffer- and TXS buffer-soluble protein fraction in comparison to vehicle-treated mice. Mean + SEM, unpaired student’s *t* test, n=6.

## DISCUSSION

α-Syn is increasingly considered a valid experimental therapeutic target for PD, based on clinical genetic and neuropathological evidence, as well as animal and cell culture studies. *SNCA* gene mutations or amplifications resulting in α-Syn pathology are tightly linked to PD pathogenesis. Moreover, α-Syn is a major constituent of Lewy-like structures, the pathological hallmark of PD and related synucleinopathies. Thus, targeting α-Syn has been a major thrust in the pharmaceutical realm. One aspect of α-Syn pathology that has been difficult to overcome is the notion that different states of its post-translational modification or aggregation differentially affect disease pathogenesis: a clear image has yet to emerge as to the real culprit of α-Syn toxicity. Although novel strategies such as anti-sense oligonucleotides, immunotherapy and small molecule inhibitors of α-Syn aggregation are being explored (45), finding a target that can be pharmacologically inhibited still holds potential as a minimally invasive and simple strategy to lower α-Syn levels – especially when such treatment course would be made over several decades. Indeed, a growing body of evidence suggests that α-Syn may play a role not only at the presynaptic space but also in the immune system (37,46,47). Therefore, careful titration of its levels may be clinically crucial. As a result, we asked whether candidates that are more amenable to traditional pharmacology (i.e. kinases) could regulate α-Syn dosage, irrespective of its aggregation status. Our previous studies identified a handful of these modifiers including TRIM28 and DCLK1 (8,22,48,49). Here, we study a heretofore unexplored target as well as its newly developed cognate inhibitor (26) for disease modification in pre-clinical models of PD: CDK14.

We show that the reduction of CDK14 protein levels is well tolerated and causes a reduction in pathogenic α-Syn accumulation in murine and human models of synucleinopathy. Genetic suppression of *Cdk14* reduces pS129 α-Syn pathology in the cortex of PFF-injected mice and dampens the development of grip strength deficits in these mice; while this rescue is not found at sites proximal to the injection, including the nigrostriatal tract. Importantly, the genetic reduction of CDK14 in DaNs derived from an individual with synucleinopathy shows equal promise in preventing phenotypic development. We show that the selective covalent CDK14 inhibitor, FMF- 04-159-2, decreases α-Syn levels in hESC-derived human neurons and mitigates PFF-induced α-Syn pathology in rat cortical neurons. Lastly, we demonstrate that administering FMF-04-159-2 *in vivo* reduces α-Syn dosage and, consequently, decreases pathogenic forms of α-Syn in a humanized mouse line expressing PD-linked A53T *SNCA*. Collectively, these results show that CDK14 is a pharmacologically tractable target for synucleinopathy.

We observed that loss of Cdk14 in PFF-treated mice reduced the level of α-Syn histopathology in cortical areas, such as the somatomotor cortex. Surprisingly, we did not detect changes in the load of pS129 α-Syn-positive cells by *Cdk14* ablation in the striatum, the PFF injection site (**Fig. 1C**). In line with this, we noticed a similar PFF-induced degeneration of the dopaminergic nigrostriatal system in WT, *Cdk14*^+/-^ and *Cdk14*^-/-^ mice (**Fig. 1D** and **Fig. S2D**). These observations imply that loss of CDK14 reduces the degree of intercellular α-Syn spreading rather than protecting neurons which are directly exposed to PFFs. Indeed, when we tested this in a cultured neuron system, we found that genetic reduction of Cdk14 dramatically decreased the spreading capacity of seed-competent α-Syn (**Fig. 2**). Therefore, it is plausible that CDK14 facilitates the cell-to-cell spread of α-Syn. Genetic reduction of CDK14 using a CRISPR/Cas9- mediated strategy in stem cell-derived human neurons carrying the PD-linked *SNCA* A53T mutation lowered levels of pS129 α-Syn (**Fig. 3**), indicating that ablation of both, murine and human *Cdk14*/*CDK14* mitigates the neuronal load of this pathology-linked form of α-Syn.

Given the lack of phenotypes in the *Cdk14^-/-^* or *Cdk14^+/-^* mice, no evidence for loss of function intolerance in humans (probability of loss of function intolerance [pLI] = 0; gnomAD database, *CDK14* | gnomAD v2.1.1), (50)) and the availability of a recently developed highly selective CDK14 inhibitor, we further explored the pharmacological tractability of CDK14 in the context of synucleinopathy. Treatment of hESC-derived cortical neurons with the newly developed CDK14 inhibitor FMF-04-159-2 induced a pronounced reduction of the total α-Syn concentration, as measured by ELISA quantification (**Fig. 4A)**. FMF-04-159-2 was recently designed to provide an improved pharmacological tool for the inhibition of CDK14 as treatment for colorectal cancer (26). Interestingly, FMF-04-159-2 was described to covalently bind and inhibit CDK14 at ∼100 nM (IC_50_ = 86 nM (26)), a dosage which lowered α-Syn levels to ∼12 % of vehicle-treated controls in our *in vitro* experiments. Applying the CDK14 inhibitor to PFF-challenged rat cortical neurons reduced the amount of aggregated (Urea buffer-soluble) α-Syn species (**Fig. 4B**), phenocopying the low degree of pS129 pathology in cortical neurons of PFF-treated *Cdk14*^-/-^ mice (**Fig. 1C**) or cultures (**Fig. 2** and **Fig. S2B**). Interestingly, we found that Cdk14 accumulated in the urea-soluble fraction, specifically upon PFF treatment. To our knowledge, it is not known whether CDK14 is present in LBs or LNs in brains of PD patients. Future histopathology experiments may generate deeper insights on if CDK14 aggregates together with α-Syn in patients with synucleinopathies. Comparable to CDK14 inhibitor-treated rat neurons with PFFs, we observed a reduction of total α-Syn in the more insoluble, TXS buffer-soluble protein fraction (and in the TSS buffer-soluble fraction) of *PAC α-Syn*^A53T^ *TG* mice which received FMF-04-159-2 via intracerebral injection (**Fig. 5A** and **B**). Reduction of total α-Syn was accompanied by lower amounts of CTT α-Syn, indicating that this form of α-Syn, which increases α-Syn’s propensity to aggregate and enhances its cytotoxic effects (51–53), is modulated by CDK14. Levels of pS129 α-Syn were reduced in the more insoluble TXS fraction of CDK14 inhibitor-treated mice, again pointing towards lower degrees of aggregated, pathology-relevant forms of α-Syn. In contrast to these findings, we found paradoxically increased amounts of pS129 α-Syn in the TSS buffer-soluble fraction (highly soluble) of CDK14 inhibitor-treated mice, suggesting that Cdk14 blockage increased pS129 α-Syn in the cytosol of this α-Syn-humanized mouse line. Notably, we did not observe any macroscopic signs of PD/neuropathy-linked behavioral abnormalities in *PAC α-Syn*^A53T^ *TG* mice upon CDK14 inhibitor treatment, such as loss of motor activity or imbalance during locomotion (**Fig. S3A**), implying that higher amounts of cytosolic pS129 α-Syn do not substantially promote cerebral and functional impairments. Elevated levels of cytosolic, non-aggregated pS129 α-Syn seem to be indicative of earlier stages of synucleinopathies, as cytoplasmic networks positive for pS129 α- Syn are more commonly observed in neurons without LBs from patients with early-stage disease, than in LB-containing neurons of patients with advanced PD (54). Based on these *in vivo* CDK14 inhibitor administration experiments, we hypothesize that blockage of CDK14 activity reduces loads of pathology-linked forms of insoluble α-Syn, potentially shifting synucleinopathy progression to an earlier phase of disease development.

Our experiments provide insights in the modulatory effects on α-Syn pathology by reduced CDK14 activity in neurons from multiple mammalian species. However, we did not observe any direct interaction between both proteins, CDK14 and α-Syn. Moreover, we could not detect any kinase activity of recombinant CDK14 toward wild-type α-Syn *in vitro* (data not shown). We therefore hypothesize that loss or inhibition of CDK14 causes a decrease in α-Syn through unknown mediators, ultimately regulating α-Syn protein levels and intercellular spreading of α- Syn pathology. Future studies will help refine the mechanism whereby CDK14 regulates α-Syn.

Our results examining the genetic and pharmacological reduction of CDK14 in PD models set the stage for future pre-clinical studies. In all behavioral experiments conducted, *Cdk14^-/-^* mice were indistinguishable from WT mice (**Fig. 1B** and **Fig. S2C**). Furthermore, loss of Cdk14 did not alter the architecture of brain tissue or peripheral organs (**Fig. S1C** to **Fig. S1E**) implying that *Cdk14* loss is not deleterious *in vivo*. This is supported by human genetics where the loss of *CDK14* appears to be well tolerated. Additionally, Cdk14 inhibition *in vivo* did not induce any signs of discomfort (**Fig. S3A**), implying that pharmacological targeting of CDK14 is safe. Similarly, systemic administration of FMF-04-159-2 at a ∼140-fold higher delivery rate (50 mg/kg/day) appears to be well tolerated by mice as shown in a recent study (55) where Cdk14 inhibition mitigated the growth of lung tumors. In our study, the activity of the CDK14 inhibitor in human neurons appears to be high, affecting CDK14 (and thus α-Syn metabolism) in the nanomolar range (**Fig 4A**). Further preclinical experiments will test whether the drug rescues PD-like neuron loss and behavioral phenotypes in models of synucleinopathies; potentially paving the way for its use in humans.

In sum, we show that CDK14 inhibition causes a decrease of total α-Syn concentrations under both *ex vivo* and *in vivo* conditions, ameliorates the levels of pathology-relevant forms of α- Syn and potentially reduces cell-to-cell transmission of α-Syn. Given the strong evidence linking α-Syn levels to PD pathogenesis, we conclude that targeting CDK14 function holds promise as a potentially disease-modifying approach to treat PD.

## CONCLUSIONS

Elevated α-Syn levels are closely linked to PD. In this study, we explore the effect of inhibiting CDK14 as a pharmacologically tractable approach to decrease α-Syn levels in cultured neurons and animal models. We show that the genetic reduction of Cdk14 mitigates grip strength impairment and ameliorates cortical α-Syn pathology in PFF-treated mice, without affecting nigrostriatal pathology proximal to the injection site; Cdk14 likely acts to regulate the intercellular spread of seed-competent α-Syn. Similarly, *CDK14* ablation reduces α-Syn pathology in human dopaminergic neurons derived from PD patients. Finally, pharmacological targeting of CDK14 lowers pathological α-Syn in cultured neurons and modifies pathogenic forms of α-Syn in mice expressing PD-linked human A53T *SNCA*. Taken together, we propose CDK14 inhibition as a novel pre-clinical strategy to treat synucleinopathy.

## LIST OF ABBREVIATIONS

α-Syn: α-Synuclein
BDW: bodyweight
CDK14: cyclin-dependent kinase 14
CL: contralateral
CTT: C-terminally truncated α-Syn
DAB: diaminobenzidine
DaNs: dopaminergic neurons
DIV: days *in vitro*
ELISA: enzyme-linked immunosorbent essay
H&E: hematoxylin and eosin
hESC: human embryonic stem cell
high exp.: high exposure
hiPSC: human induced pluripotent stem cell
IL: ipsilateral
min: minutes
KO: knockout
LC-MS/MS: liquid chromatography – mass spectrometry/mass spectrometry
Mono: α-Synuclein monomers
NPC: neural precursor cell
PBS: phosphate-buffered saline
PD: Parkinson’s disease
PFFs: α-Synuclein preformed fibrils
RT: room temperature
sec: seconds
SEM: standard error of the mean
SN: *substantia nigra pars compacta*
TG: transgene
UT: untreated
WT: Wildtype

## DECLARATIONS

### Ethics approval and consent to participate

Animal experiments were done under the approved breeding and behavior protocols approved by the University of Ottawa Animal Care Committee. Studies with hESCs were performed following approval by the Stem Cell Oversight Committee of Canada and the Institutional Review Board (Ottawa Health Science Network Research Ethics Board).

### Consent for publication

Not applicable

### Availability of data and material

All data of this study are in the main text or in the Supplementary Materials. Cdk14^-/-^ mice were obtained from David S. Park from the University of Calgary and are on an F14 C57BL/6N background. *PAC α-Syn*^A53T^ *TG* mice (dbl-PAC-Tg(*SNCA*^A53T^)^+/+^;*Snca*^-/-^, (28)) were provided by Robert L. Nussbaum from the University of California, San Francisco. Any other materials are commercially available.

### Funding

Parkinson’s Foundation-APDA Summer Student Fellowship PF-APDA-SFW-1919, Parkinson Research Consortium (PRC) Summer Studentship, University of Ottawa Faculty of Medicine, Office of Francophones Affairs Scholarship, Undergraduate Research Scholarship (URS), University of Ottawa. (JLAP)

Parkinson Research Consortium (PRC) Larry Haffner Fellowship, Parkinson Canada Basic Research Fellowship BRF-2021-0000000048. (KMR)

Parkinson’s Research Consortium (PRC) Bonnie and Don Poole Fellowship, Ontario Graduate Scholarship, Queen Elizabeth II Scholarship. (HMG)

Canadian Institutes of Health Research (FRN-153188). (WLS) Vanier Canada. (MGS)

Parkinson’s Society of Southwestern Ontario (PSSO). (BBD) Canadian Institutes of Health Research (FRN-159443). (SDR)

Parkinson’s Foundation Stanley Fahn Junior Faculty Award (PF-JFA-1762), Canadian Institutes of Health Research (PJT-169097), the Parkinson Canada New Investigator Award (2018-00016), Aligning Science Across Parkinson’s (ASAP-020625) through the Michael J. Fox Foundation for Parkinson’s Research (MJFF). (MWCR)

### Authors’ contributions

Conceptualization: JLAP, KMR, MWCR

Methodology: JLAP, KMR, MS, BBD, EL, BN, NAL, HMG, AB, MGS, BBD, SMC, MWCR

Investigation: JLAP, KMR, MS, BBD, HMG, SMC, MWCR

Supervision: WLS, MGS, SDR, MWCR

Writing – original draft: JLAP, KMR, MWCR

Writing – review & editing: JLAP, KMR, JJT, MGS, WLS, PB, SDR, MWCR

Resources: AJ, JM, PB

## Acknowledgements

The authors thank Drs. Eliezer Masliah (UCSD/NIA), Robert L. Nussbaum (UCSF), and David Park (University of Calgary) for sharing the genetically modified mice used in this study. We thank Dr. G. Vázquez-Vélez (Washington University) for his critical appraisal of the manuscript. The authors also thank the following Core facilities from the University of Ottawa and the Ottawa Hospital Research Institute for use of their facility, equipment, and expertise: Animal Behaviour and Physiology Core (RRID: SCR_022882), Cell Biology and Imaging Acquisition Core (RRID: SCR_021845), StemCore Laboratories (RRID: SCR_012601) and Louise Pelletier Histology Core (RRID: SCR_021737). The authors also thank The Metabolomics Innovation Centre (TMIC) for the analysis of the striatal dopamine content and Anthony Carter (Surgical Core Service, University of Ottawa) for his support in stereotactic surgeries. Schemes were generated using biorender (https://biorender.com/).

