## Supplemental information for "Genetic and pharmacological reduction of CDK14 mitigates α-synuclein pathology in human neurons and in rodent models of Parkinson’s disease"

**Supplementary Materials**

**Figure S1. Cdk14 distribution in mouse organs and organ architecture of *Cdk14*$^{-/-}$ mice.** (A) Micrographs from *in situ* hybridization experiments (downloaded from the Allen Brain Atlas) depicting the expression of *Snca* and *Cdk14* in the mouse brain. SN and Hippocampus are highlighted at 10x magnification. (B) Immunoblots showing Cdk14 protein from different organs of *Cdk14*$^{-/-}$ and WT mice. (C) Hematoxylin and eosin (H&E) staining does not reveal obvious rearrangements of brain morphology of *Cdk1*$^{+/-}$ and *Cdk14*$^{-/-}$ mice in comparison to WT-mice (1 mm and 50 µm scale bars). (D) Synapsin and PSD95 immunofluorescence experiments in primary cortical neurons and in the hippocampus of adult mice do not display differences in synaptic integrity between WT, *Cdk1*$^{+/-}$ and *Cdk14*$^{-/-}$ mice (20 µm scale bar). Mean + SEM, one-way ANOVA, Bonferroni *post hoc* comparisons, n=3-4. (E) H&E-stainings of lungs and spleens of *Cdk1*$^{+/-}$ and *Cdk14*$^{-/-}$ mice do not differ from stainings of WT mice (100 µm scale bars). (F) Immunoblots showing the levels of α-Syn, Cdk14, and GAPDH in the TSS buffer-soluble protein fraction from hemibrains of 3-month-old WT, *Cdk14*$^{+/-}$ and *Cdk14*$^{-/-}$ mice. Mean + SEM, one-way ANOVA, Bonferroni *post hoc* comparisons, n=4-5.

**Figure S2. Behavioral profile and numbers of midbrain dopaminergic neurons of PFF injected *Cdk14*$^{-/-}$-mice.** (A) Transmission electron micrograph (500 nm scale bar) illustrating mouse α-Syn PFFs used for intrastriatal injections and corresponding α-Syn PFF length analysis. (B) Immunofluorescence experiment depicting less pS129 α-Syn-positive neurons in primary cortical neurons from *Cdk14*$^{-/-}$ mice than in cultured neurons from WT-mice after PFF-treatment at 9 DIV. Mean + SEM, one-way ANOVA, Bonferroni *post hoc* comparisons, n=4 (50 µm scale bar). (C) 12-month-old *Cdk14*$^{+/-}$ and *Cdk14*$^{-/-}$ mice do not display altered behavior in the nesting, tail suspension, elevated plus maze, Y maze, open field, hanging wire, pole (Mean + SEM, two-way ANOVA, Bonferroni *post hoc* comparisons) and rotarod test (Mean +/- SEM, repeated measures two-way ANOVA, Bonferroni *post hoc* comparisons, n:9-10). PFF treatment does not result in significant changes of behavior at 6 months post injection. (D) PFF injection does not change the striatal dopamine content in the ipsilateral (IL), PFF-injected hemisphere (measured by LC-MS/MS, mean + SEM, two-way ANOVA, Bonferroni *post hoc* comparisons, n:2-8), but it decreases the number of TH-positive cells in the *substantia nigra pars compacta* (SN) IL relative to the non-injected, contralateral hemisphere (CL) to a similar extent in WT-, *Cdk14*$^{+/-}$- and *Cdk14*$^{-/-}$- mice at -3.08 mm relative to bregma (sections counterstained with hematoxylin, 500 µm scale bar). Mean + SEM, two-way ANOVA, Bonferroni *post hoc* comparisons, n=3. (E) Immunoblots showing the levels of α-Syn, Cdk14, and GAPDH in the TSS buffer-soluble protein fraction from hemibrains of 3-month-old WT, *Cdk14*$^{+/-}$ and *Cdk14*$^{-/-}$ mice. Mean + SEM, one-way ANOVA, Bonferroni *post hoc* comparisons, n=4-5.

**Figure S3. *In vivo* administration of FMF-04-159-2 does not induce pain or alter organ morphology.** (A) The development of body weight (BDW, as % relative to the day of surgery (day 0)), the activity, neurological signs for pain, the facial grimace, the coat condition and the respiration (scored from 0 to 3, 3 = maximal discomfort) of FMF-04-159-2-treated *PAC α-Syn*$^{A53T}$ *TG* mice compared to their vehicle-treated counterparts were measured over a period of 25 days after the stereotactic surgery. FMF-04-159-2 was applied at 0.35 mg/kg/day for 28 days in 4-month-old *PAC α-Syn*$^{A53T}$ *TG* mice. Mean +/- SEM, repeated measures two-way ANOVA, Bonferroni *post hoc* comparisons, n=6. (B) H&E staining of the lung, spleen and liver of FMF-04-

159-2-treated *PAC α-Syn*$^{A53T}$ *TG* mice did not reveal alterations in organ cytoarchitecture compared to vehicle-treated mice (100 µm scale bars).

**Fig. S1.**

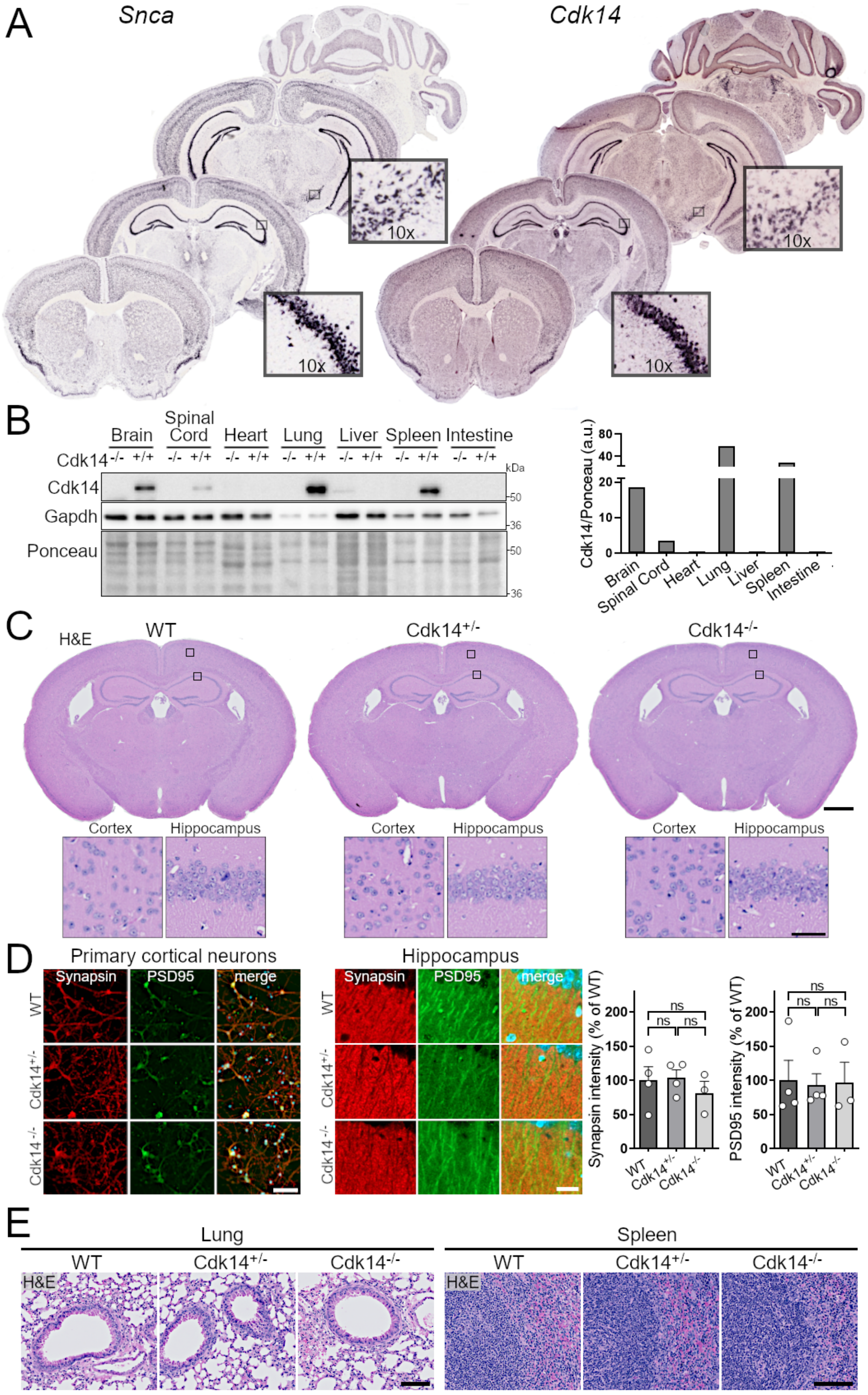
A
Snca
Cdk14
10x
10x
10x
10x
B
Brain
Spinal Cord
Heart
Lung
Liver
Spleen
Intestine
Cdk14 -/- +/+ -/- +/+ -/- +/+ -/- +/+ -/- +/+ -/- +/+ -/- +/+
kDa
Cdk14
Gapdh
Ponceau
50
36
50
36
Cdk14/Ponceau (a.u.)
80
40
20
10
0
Brain
Spinal Cord
Heart
Lung
Liver
Spleen
Intestine
C
H&E
WT
Cdk14+/-
Cdk14-/-
Cortex
Hippocampus
D
Primary cortical neurons
Hippocampus
Synapsin
PSD95
merge
WT
Cdk14+/-
Cdk14-/-
Synapsin intensity (% of WT)
PSD95 intensity (% of WT)
200
150
100
50
0
ns
E
Lung
Spleen
WT
Cdk14+/-
Cdk14-/-
H&E

1103 **Fig. S2**

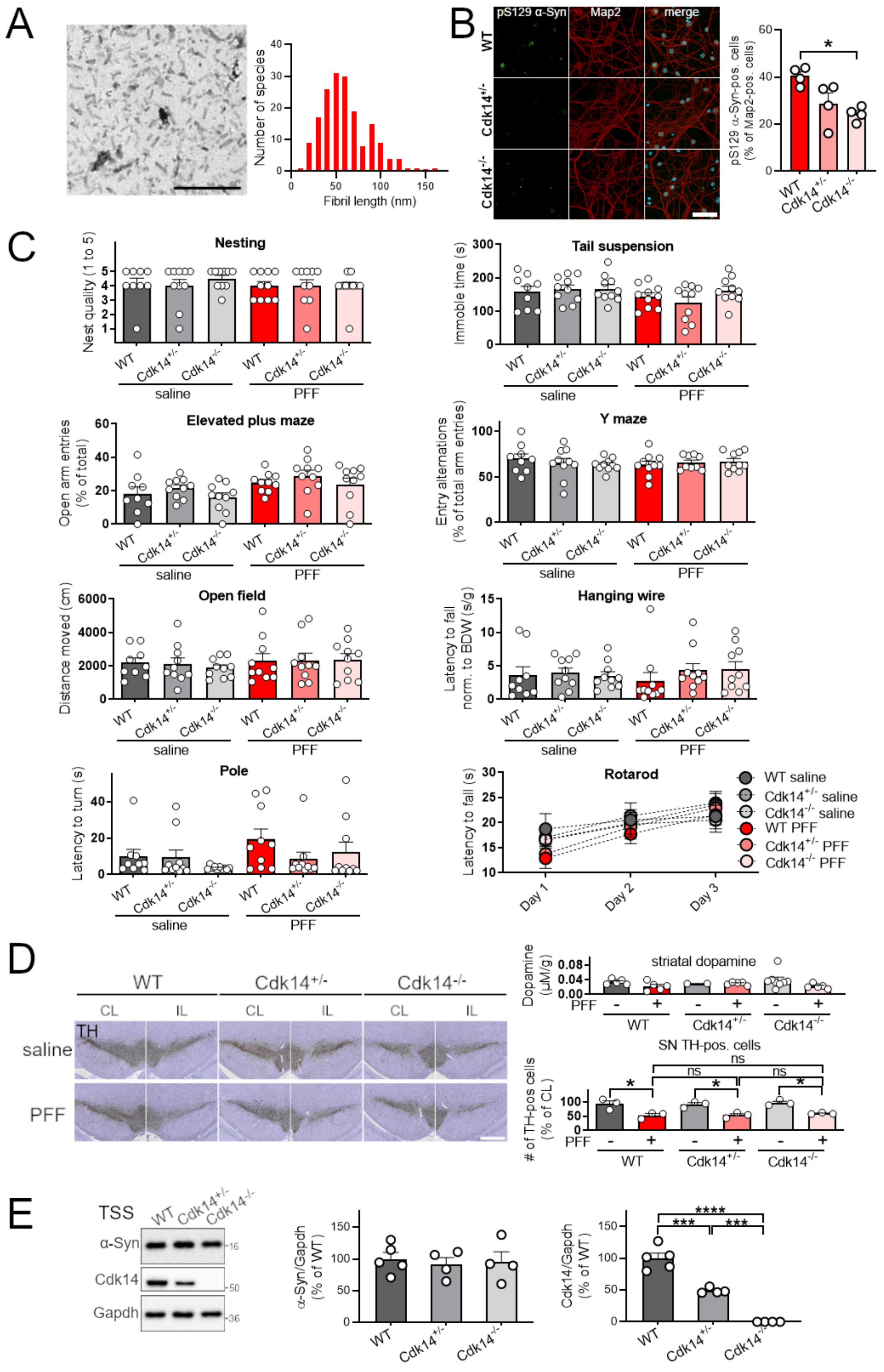


1104

1105 **Fig. S3**

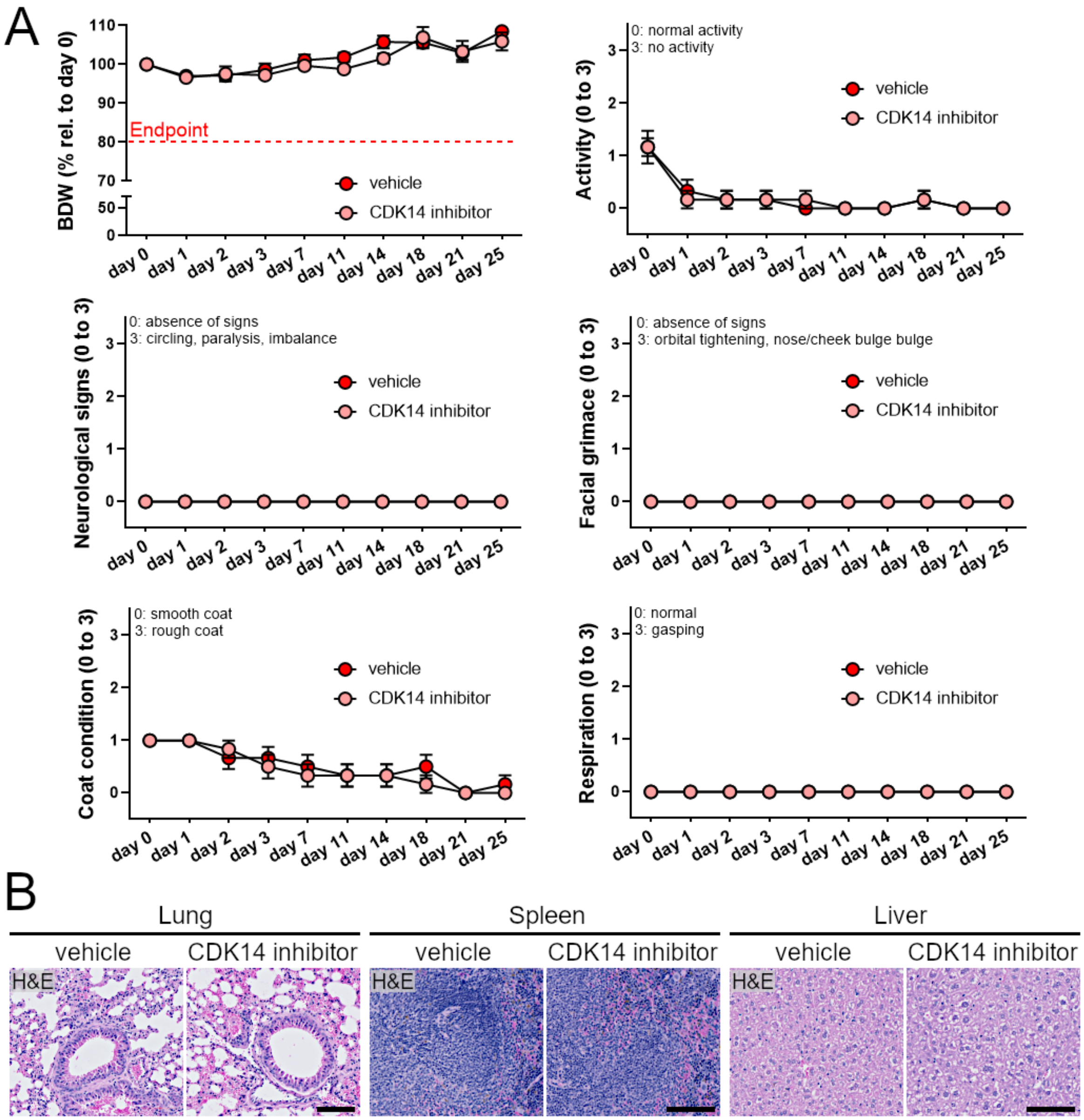


1106
